# Factors contributing to the accumulation of reproductive isolation: a mixed model approach

**DOI:** 10.1101/072264

**Authors:** Dean M. Castillo

**Author notes:** Current Address: Department of Molecular Biology and Genetics, 526 Campus Road, Cornell University, Ithaca, New York 14853.

## Abstract

The analysis of large data sets describing reproductive isolation between species that vary in their degrees of relatedness has been extremely influential in the study of speciation. However, several limitations make it difficult to test specific hypotheses about which factors predict the evolution of reproductive isolation. In particular, the statistical methods typically used are limited in their ability to test complex hypotheses involving multiple predictor variables or interactions between variables; at least one method, the Mantel Test, has also been found to be unreliable. In this paper, I describe a framework to determine which factors contribute to the evolution of reproductive isolation using phylogenetic linear mixed models. Phylogenetic linear mixed models do not suffer from the same statistical limitations as other methods and I demonstrate the flexibility of this framework to analyze data collected at different evolutionary scales, to test both categorical and continuous predictor variables, and to test the effect of multiple predictors simultaneously, all of which cannot be achieved using any other single statistical method. I do so by re-analyzing several classic data sets and explicitly testing hypotheses that had previously been untested directly, including differences in accumulation of reproductive isolation between sympatric and allopatric species pairs.

## Introduction

The divergence of lineages, in the process known as speciation, is only complete after gene flow is reduced following the evolution of reproductive isolation. The two general modes of reproductive isolation are prezygotic and postzygotic barriers to reproduction. Patterns of reproductive isolation inferred from comparative analyses of species willingness/ability to mate (i.e. prezygotic isolation) and produce viable and fertile offspring (i.e. postzygotic isolation) have generated important and influential observations about the evolution of reproductive isolation. These studies have been especially important for making generalizations about comparisons between allopatric and sympatric species pairs, or prezygotic and postzygotic barriers to reproduction (Coyne and Orr 1989; Presgraves 2002; Mendelson 2003; Moyle et al. 2004; Funk et al. 2006; Malone and Fontenot 2008; Meiners and Winkelmann 2012). These analyses have produced iconic patterns/rules of speciation including that reproductive isolation accumulates more quickly between sympatric species pairs compared to allopatric species pairs and, specifically, that in sympatry prezygotic reproductive isolation appears more quickly than postzygotic reproductive isolation. Nonetheless, the contemporary methods used to analyze these data are known to suffer from both statistical errors and analytical limitations that prevent a more robust simultaneous assessment of the factors that most strongly influence the accumulation of reproductive isolation.

The two methods most typically used to analyze comparative data on reproductive isolation (hereafter, crossability) are Mantel tests, including partial mantel tests (Mantel 1967; Smouse et al. 1986), and phylogenetic regression based on independent contrasts (PICs; Felsenstein 1985; Grafen 1989). These two methods (Mantel/matrix regression vs. PICs) reflect differences in the scale of relationships between lineages that are being tested. Studies that use PICs, or node based averages (Fitzpatrick and Turelli 2006), generally have some information about phylogenetic or exact sister species relationships, which is needed to accurately calculate independent contrasts. The use of PICs, or methods that assume independence of lineages, are not appropriate for intraspecies data (Slatkin et al. 2002; Stone et al. 2011). Thus, Mantel tests are applied when specific phylogenetic relationships are unknown (if few molecular markers are used) and/or lineages being tested include extensive sampling from only a few closely related species. The main limitations of these approaches are threefold: lack of statistical power, failure to deal adequately with non-independence, and the inability to test categorical variables and/or multiple variables simultaneously. The first two limitations have been discussed in detail for the Mantel test and it has been determined that Mantel tests can have unacceptably high type-I error rates (Legendre 2000; Harmon and Glor 2010). The last limitation results in the inability to test biologically interesting hypotheses, and applies equally to both Mantel test and PIC type tests. For example, in many of the classic studies the comparisons between sympatric vs. allopatric rates of evolution were never formally tested with a statistical model. Recent attempts to refine the patterns of the accumulation of reproductive isolation have also included other explanatory variables, including ecological differences, range size overlap, and other traits (Yukilevich 2012; Turelli et al. 2014), but have not tested multiple variables simultaneously or the interaction between these variables.

Categorical variables are of particular interest to most researchers as they can describe geographic relationships (allopatry vs. sympatry), mating systems (animals: monogamy vs. polygamy, plants: outcrossing vs. selfing), or phenotypic traits (e.g., pigmentation/patterning) that could have very strong influence on the rate and strength of isolation accumulation. Categorical variables can be difficult to model in Mantel tests because they rely on pairwise distance matrices that require no missing data (Mantel 1967). When categorical variables can be represented as distances, multiple matrix regression can be used for analysis (Legendre and Fortin 2010; Wang 2013). This is typically only occurs with binary categorical variables because creating predictor variables based on distances for categorical variables with multiple levels becomes unfeasable. Therefore, Mantel tests based on pairwise distance cannot accommodate hypotheses that test differences between levels of categorical variables.

Regression models are the most promising for incorporating categorical variables, but the analysis is not simple when the regression is carried out using PICs. Continuous variables can be accommodated in PIC models because they are assumed to evolve under Brownian motion; this is what enables node values to be estimated and contrasts to be evaluated. However an analogous model does not exist for categorical variables, unless all daughter taxa (taxa derived from a common ancestor) share the same categorical trait value (Burt 1989). Even with this conservative approach the number of contrasts would be reduced tremendously in studies of crossability, and may not be applicable to most systems. One way of circumventing this issue would be to analyze the cross product of a categorical and continuous variable (Garland et al. 1992). Although this method would take the model a step further by analyzing differences in slope (assuming genetic distance is a covariate), it cannot be used to estimate mean crossability because regression in PICs is constrained to pass through the origin.

To accommodate more complex hypotheses, a more appropriate and powerful analytical framework should be: 1) flexible to test explicit hypothesis with multiple variables, both continuous and categorical, and 2) able to handle different data types to cover differences in taxonomic scale. Here I describe a framework that overcomes several limitations of current approaches to analyzing these data. Specifically, this framework can be used to test whether geographical context, and/or any other trait hypothesized to be important to speciation, contribute to patterns of crossability using comparative crossing data and phylogenetic linear mixed effect models (similar to phylogenetic least squares, PGLS). Using linear models allows flexibility to test many different hypotheses and can include multiple predictors, and their interactions simultaneously. The advantage of this framework over PICs is that the phylogenetic structure is modeled as a covariance matrix, and thus contrasts for categorical predictors do not have to be calculated. The use of this covariance matrix allows/accounts for phylogeny by either using a pairwise distance matrix or a phylogeny. I use this framework to reanalyze classic data sets (*Drosophila*, Bufonidae, *Silene*) making explicit statistical comparisons between allopatric and sympatric conditions as an example of how categorical variables can be included in these types of analyses. In addition, I test two new hypotheses about the relationship between floral differences and crossability in each of the plant genera *Silene* and *Nolana.*

## Methods

### Description of Linear Mixed Model Approach

In their most basic form, linear mixed models include a response variable (*y*), fixed effects (*XB*), random effects (*Zu*), and residuals (*e*), where *X* is the fixed effect design matrix, *B* is the vector of estimated coefficients, *Z* is the random effect design matrix, and *u* is a vector of random effect estimators.

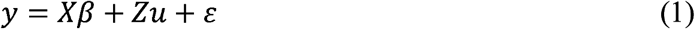

Relatedness either via pedigree or phylogeny is incorporated into a structured matrix (typically *A*, the additive genetic relatedness matrix, see (Hadfield 2010; Hadfield and Nakagawa 2010) that is considered known and contributes to defining the variance structure of random effects in the model. In equation (2) the random effect estimators are distributed normally with mean=0 and variance matrix *G.* This matrix can be decomposed into a standard variance/covariance matrix *V*, and the structured matrix *A* (Equation 3)

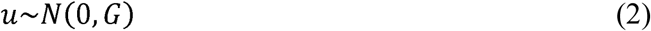

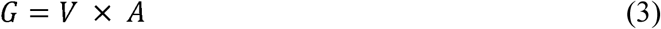

This modeling framework can also accommodate a pair-wise genetic distance matrix (*D*) because of its similarities to the additive genetic relatedness matrix. Each element of *A*, in the absence of inbreeding, represents the expected proportion of genes shared by two individuals. Values range from 0 (completely unrelated individuals) to 1 (completely identical individuals). Whereas some measures of genetic distance are not bound by 0 and 1, (Euclidean distances, Nei's standard genetic distance (Nei 1972)), others formally meet this requirement (Nei's D_A_ distance (Nei et al. 1983), Wier and Cockerham *θ* (Weir and Cockerham 1984)), and in most actual studies values are rarely seen above 1. Genetic distance values of 0 represent more closely related populations. To make the genetic distance matrix analogous to the structured *A* matrix I use 1-*D* in my analyses so that smaller values (closer to 0) represent more distantly related species. Genetic distance used as a predictor variable remained unmanipulated.

The datasets I analyze include *Drosophila* (Coyne and Orr 1989; Yukilevich 2012; Nosil 2013; Turelli et al. 2014), Bufonidae (Malone and Fontenot 2008), *Silene* (Moyle et al. 2004), and *Nolana* (Jewell et al. 2012). All of these datasets have been analyzed previously for patterns of crossability and are explained in detail below.

### Model Descriptions

The phylogenetic mixed model framework can be applied to any case where the goal is to test the effect of a categorical variable and/or multiple variables on the evolution reproductive isolation. I illustrate this with several examples. In one set of analyses I estimate the effects of a continuous trait (genetic distance) and a categorical trait (allopatry vs. sympatry; or trait presence vs. absence) and their interaction. In another analysis, I analyze three continuous variables simultaneously (genetic distance, geographic distance, and a phenotypic/morphological distance).

#### Model incorporating relatedness and categorical biogeography

A specific model that can be analyzed to determine if reproductive isolation is different in crosses between sympatric species compared to allopatric species, and how this relationship changes with time since divergence, will incorporate the crossability data (the measure of reproductive isolation, either pre-zygotic isolation, post-zygotic isolation, or total isolation) as a function of genetic distance and geographic context (sympatry vs. allopatry) while controlling for phylogenetic relatedness (Eq. 4).

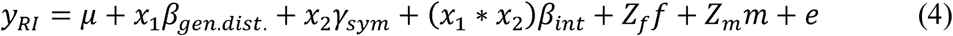

The first term (*µ*) is the intercept of the linear model, and can be thought of as the baseline level of reproductive isolation if there is no significant relationship between reproductive isolation and genetic distance. If there is a significant relationship the intercept then represents the reproductive isolation between closely related species. The variable *x_1_* is a vector of genetic distance between the species pair and (*β_gen.dist._* is the slope of the relationship between reproductive isolation and genetic distance. The variable *x_2_* is a dummy variable (0 when the species pair is allopatric and 1 when the species pair is sympatric) and *γ_sym_* is the potential difference in reproductive isolation that can be attributed to the species occurring in sympatry. (*β_int_* is the potential change in the slope of the relationship between reproductive isolation and genetic distance for sympatric species crosses. Lastly there are two *Z* matrices; they represent the identity of the female parent species and male parent species used in the cross (or can be designated species 1 and species 2 if sex of the parents is not important). I use a separate effect for each parent in the cross because not all species are used as both a male parent and female parent, depending on the data set. Recent studies use interactions between phylogenetic effects for species interactions (Hadfield et al. 2014), but for the datasets in this study there are very crosses made outside of very closely related species so any matrix describing an interaction between phylogenetic effects would be very sparse and hard to estimate. Using this model I was able to test for differences in the rate of reproductive isolation for three data sets: *Drosophila*, Bufonidae, and *Silene* (described below). The *Drosophila* data set has previously been shown to have increased levels of prezygotic reproductive isolation between sympatric pairs compared to allopatric pairs (Coyne and Orr 1989), but these rates were never directly compared in a single model.

#### Model incorporating relatedness and categorical trait differences

As a second example, I incorporate a categorical variable with more than two levels, such as floral color differences/similarities between the species pair, by using the same model described above but with flower color substituted for geographic context.

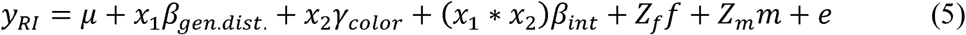

Here the *x_2_* variable can have two levels, for example, *x_2_* will have two levels if one level represents crosses between species that share the same state for floral color (red × red and white × white) and the other level represents crosses between species with different floral colors (red × white and the reciprocal). Alternatively the *x_2_* variable could have multiple levels: crosses between species that both have red flowers, crosses between species that both have white flowers, and crosses between red and white flowered species. This model would test whether specific floral colors/morphologies have increased speciation rates compared to other floral morphologies.

Using the *Silene* dataset I test the hypothesis that floral differences may contribute to post-mating and postzygotic isolation. Flower color and shape (together making up a pollination syndrome; Fenster et al. 2004) are often thought to contribute to premating reproductive isolation through pollinator preference (Schemske and Bradshaw 1999; Kay and Sargent 2009) and mechanical isolation, including pollen placement (Grant 1992; Hodges and Arnold 1994; Smith and Rausher 2007). Additionally pollination syndromes can contribute to post-mating isolation if there are differences in the ability of pollen to reach the ovary depending on differences in style length (Lee et al. 2008). Moreover, genes that control floral development are active throughout several developmental processes including gametogenesis and embryonic development (Smaczniak et al. 2012), so that post-zygotic isolation may evolve as a byproduct of floral divergence (Haak et al. 2014).

#### Model incorporating multiple continuous variables with potential correlations

Often geographical context may not be captured by a categorical variable (allopatry vs. sympatry) but instead by a continuous variable such as geographic distance. When spatial variation is included in analyses we expect correlations between geographic/spatial variables and other variables of interest including genetic distance. We expect that lineages that are more geographically isolated are also more genetically differentiated. Additional correlations may exist between morphological traits that can contribute to reproductive isolation, such as quantitative floral traits. This model allows multiple continuous variables to be analyzed simultaneously and accounts for correlations between the predictor variables (Equation 5). In the example illustrated below, I include genetic distance, geographic distance, and the difference in corolla tube length, which is one measure that captures the difference in floral size between species.

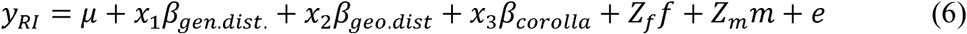

If these variables are correlated with one another it will be difficult to make inferences on any given parameter estimate since mulitcollinearty can cause variance inflation (increased estimates of variance parameters compared to model where variables are not correlated). To address this issue I explicitly allow for covariance between these variables in the model by changing the variance structure from being independent (Equation 6), to being correlated (Equation 7).

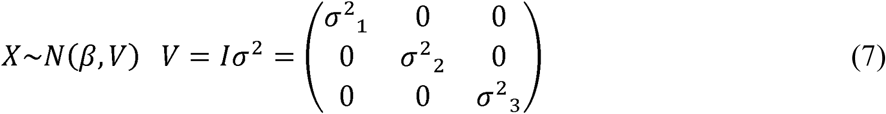

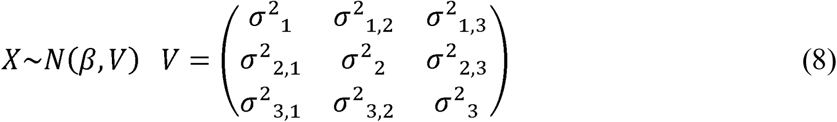

This model can be applied to the *Nolana* dataset, for example, to disentangle the effects of multiple correlated variables. In the original analysis of the *Nolana* dataset (Jewell et al. 2012) there was some evidence that genetic distance and/or geographic distance predicted reproductive isolation. However, since these two variables were highly correlated that analysis lacked power to disentangle this pattern using partial mantel tests. Using the approach here, I directly modeled the correlation between the continuous variables by including a covariance/correlation matrix between the three continuous variables.

#### Model incorporating genetic distance matrix in place of phylogeny

Another way to correct for relatedness in this framework is to use a pairwise genetic distance matrix. This method would be most suitable for closely related species for which constructing a phylogenetic tree is inappropriate. As an example, I analyzed the *Nolana* data using information from a pairwise genetic distance matric (specifically 1-*D*, where *D* is the genetic distance matrix) in the place of the structured *A* matrix, to see if the inferences varied between the two methods. The package MCMCglmm (Hadfield 2010) requires the inverse of the structured matrix and has built in functions to take the inverse of the relatedness matrix from either pedigree or phylogeny information. To use the genetic distance matrix, I used the ginv function from the MASS (Venables and Ripley 2002) package to find the generalized inverse of the 1-*D* matrix.

### Interpreting model outputs

In the Bayesian framework used in MCMCglmm, there is no formal distinction between fixed and random effects, but for ease of explanation I will call all parameters that are estimated for the predictor variables fixed effects, and the phylogenetic variance random effects. I can test any specific hypothesis for fixed effects by examining the highest posterior density (HPD) of the parameter from a given model. For example, to test the hypothesis that there is a difference in average reproductive isolation between allopatric and sympatric species pairs, I determine if the HPD for regression coefficient (*γ_sym_*) includes 0. If *γ_sym_* does not include 0 than there is evidence supporting the hypothesis of a significant difference in reproductive isolation between allopatric and sympatric species pairs. Similarly, testing the hypothesis that the rate of accumulation of reproductive isolation differs between two groups (e.g. sympatric and allopatric pairs), involves evaluating the slope *β_int_* to again assess whether the HPD includes 0. For all analyses I set the prior distributions for random effect covariances and residuals as follows. MCMCglmm uses inverse Wishart distributions for random effects, with scale parameter *V*, and degrees of freedom parameter *n*. For each model I set *V* to be 1/3 the total variance in the response variable because there were three covariance matrices (two matrices for male and female parent, and the residual covariance matrix).

For each model I ran two MCMC chains; this enabled me to determine lack of convergence by examining within and between chain properties. The trace plots (iteration number vs. value of the draw) were visually inspected for all variables/chains, to see if chains were mixing well or if there was high auto-correlation (which would signify chain being stuck on a local maximum and thereby give a false signal of convergence). Along with visual inspection, the primary tool used to determine if the MCMC chains failed to converge was the Gelman-Rubin Diagnostic (Gelman and Rubin 1992). This test uses information from the variance of the mixture of both chains, and the variance within a single chain to calculate a potential reduction factor (Gelman and Rubin 1992; Brooks and Gelman 1998). A value of 1 indicates that the chains have converged because the ratio of variance between and within chains is identical. In practice chains are typically run until the reduction factor for all variables is less than 1.1 (Brooks and Gelman 1998). I used the coda package (Plummer et al. 2006) in R to calculate the convergence diagnostics. I considered a model to have converged if the all scale reduction factors for all variables (both fixed and random effects) were less than or equal to 1.1 and the trace plots indicate good mixing (actual scale reduction factors were generally less than 1.06, and the multivariate scale reduction factors were less than 1.02.) Code for all analyses are available through Dryad digital repository.

### Data Sets

Four different data sets were used here and are explained in detail below; as outlined, in several cases these were enriched with additional tree construction, biogeographical information, and/or trait data prior to performing analyses. All data sets and phylogenies are available Dryad digital repository.

#### Drosophila

The *Drosophila* dataset is an expansion of the original data set used by Coyne and Orr (Coyne and Orr 1989) and was accessed from http://www.drosophila-speciation-patterns.com/. In this dataset, prezygotic isolation estimates are based on choice and no-choice mating assays, depending on the specific species pair. Post-zygotic isolation is a combination of hybrid sterility and inviability of any hybrid progeny that were produced. To include phylogenetic information in my model, I combined data from two phylogenies that had complementary information and largely agreed on phylogenetic relationships. The first phylogeny (van der Linde et al. 2010; VL) provides a useful backbone for different species groups, but lacks species richness within some groups. The second phylogeny (Morales-Hojas and Vieira 2012; MH) includes more species for particular clades. Since the VL tree includes more species groups in total I used this tree to define relationships between the major species groups. The VL tree was constructed from a supermatrix and I was able to combine it with the MH as follows. After scaling both trees so that they were ultrametric I could substitute species relationships from the MH tree into the VL tree, by transforming the branch length from the MH tree, so they were proportional to the branch length of the corresponding clade in VL tree (see example: Supplementary Fig. 1). For some groups, crossability data included races/subspecies and these were represented as polytomies. After constructing the phylogeny I only retained crossing data where both species were present in the phylogeny, leaving me with 182 crosses total to include in analyses.

#### Bufonidae

The primary dataset consisted of post-zygotic isolation estimated from in vitro crosses (Malone and Fontenot 2008). Several stages of development were used to calculate a reproductive isolation index including: fertilization rate, hatching rate, the number of tadpoles produced, percentage of tadpoles metamorphosed, fertility in backcross analysis, and the stage at which eggs ceased to develop. The reproductive isolation index was then calculated similarly to (Coyne and Orr 1989; Presgraves 2002). To evaluate the sensitivity of the inferences to this index, I also conducted analyses on an additional reproductive isolation index that takes into account that these barriers are sequential (Ramsey et al. 2003). This made the response variable more continuous instead of considering only a fixed number of values.

I used the original sequence alignment to construct a neighbor-joining tree replicating the original analysis (Malone and Fontenot 2008). In addition, I also enriched this dataset by determining allopatry/sympathy relationships, by downloading shape files (vectors storing geometric information) for each species from the IUCN Red List Database (IUCN 2014.). These files were reprojected (changed from 3D to 2D objects) to Albers Equal areas using rgdal (Bivand et al. 2013) and maptools (Bivand and Lewin-Koh 2013) in R with parameters specific for the region where they were located (Asia, Africa, Europe, or North America). I then determined whether species ranges overlapped using PBSmapping (Schnute et al. 2012) in R. Species that had no overlap were designated as allopatric.

#### Silene

The original crossing data was compiled in Moyle et al. (2004). Prezygotic isolation was a measure of the total number of failed pollinations (likely due to pollen pistil interactions) in interspecific crosses, compared to the crossability of the parental species. Postzygotic isolation was estimated from pollen sterility of F1 hybrids. The original data included sympatric and allopatric relationships and I enriched the dataset by including flower color for each species. Floral color data were summarized from the available literature, online flora projects, and personal observations. The phylogeny comes from a super tree (Jenkins and Keller 2011). Similar to the *Drosophila* data I only retained taxa for analysis that could be placed into the phylogeny, and I allowed polytomies for certain taxa where different subspecies were used in crosses. This yielded 65 crosses in total, for analyses.

#### Nolana

The *Nolana* data were originally presented in Jewell et al. (2012). Since the authors found no measurable prezygotic isolation, I focused on total post-zygotic isolation that was a combination of fruit set, mericarp size, and seed set. The phylogeny used in the original study had several large polytomies. To resolve these polytomies I used the original sequence data to construct a new phylogeny in Raxml (Stamatakis 2014), by allowing each gene (genes on chloroplast were concatenated and treated as one unit) to have its own substitution model. These data also contained a complete pairwise genetic distance matrix, pairwise measures of geographic distance, and pairwise measures of differences in specific aspects of floral morphology, all three of which are significantly correlated with one another. In my analyses I used corolla diameter differences to quantify floral distance, though similar results were achieved with another measure of floral divergence (corolla depth difference) likely because these measures were highly similar and correlated (data not shown).

## Results

### Drosophila

Using the *Drosophila* crossability data I tested the hypothesis that there are differences in the level of reproductive isolation between allopatric and sympatric species pairs (*µ*=baseline for allopatry and *γ_sym_* = change in RI for sympatry) and that reproductive isolation accumulates at a different rate between allopatric and sympatric species pairs (i.e. slope between genetic distance and RI differs; *β_gen.dist_*=relationship between genetic distance and RI for allopatric pairs and *β_int_*. = change in slope for sympatric pairs). To interpret these models it is often easiest look at which coefficients contribute to reproductive isolation in allopatric vs. sympatric species pairs. The overall model has many coefficients but only a few differentiate the allopatric vs. sympatric species pairs. The overall model has the following form:

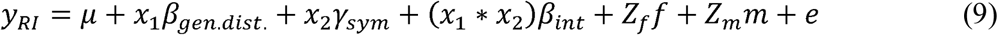

The variable *x_1_* is a continuous variable, but the variable *x_2_* is binary and is equal to 0 for allopatric pairs and 1 for sympatric pairs. Thus for allopatric pairs the model simplifies to:

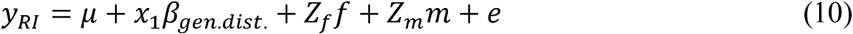

For sympatric pairs the model simplifies to

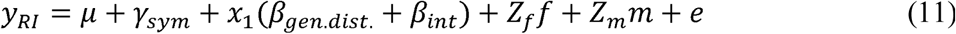

This highlights that the coefficient *β_int._* represents the change in slope for sympatric pairs and that the total slope for sympatric species pairs is the sum of the two beta coefficients.

For prezygotic isolation the intercept (baseline level of reproductive isolation) was significantly different than zero, and reproductive isolation was significantly elevated in sympatric pairs (*γ_sym_*= (0.2533, 0.4764), (Lower 95% HPD interval, Upper 95% HPD interval); Table 1). There was also a significant relationship between genetic distance and reproductive isolation, as detected in the original study (Coyne and Orr 1989), but only for allopatric species pairs. In comparison, the overall relationship between genetic distance and reproductive isolation is non-significant for sympatric pairs (Table 1). In this model the coefficients are additive (Equation 11), and the relationship between genetic distance and reproductive isolation is not significantly different from zero (HPD for (*β_gen.dist_*+ *β_int_*=(-0.1250, 0.3571)). The effect of sympatry on reproductive isolation (*γ_sym_*) is so strong that most of the reproductive isolation values are near 1 across all genetic distances (completely isolated; Fig. 1).

**Table 1.** Summary of coefficients estimated for the analysis of prezygotic (left) and postzygotic (right) reproductive isolation from the *Drosophila* data. The confidence intervals are for 95% Highest posterior density (HPD) and are significant if they do not include zero (in bold).

| Coefficient | Biological meaning | Prezygotic |  | Postzygotic |  |
| --- | --- | --- | --- | --- | --- |
|  |  | Lower | Upper | Lower | Upper |
| $\mu$ (intercept) | average RI | **0.1975** | **0.6033** | -0.0005 | 0.4001 |
| $\beta_{gen.dist.}$ | slope relating genetic distance and RI | **0.1959** | **0.4164** | **0.1906** | **0.4644** |
| $\gamma_{sym}$ | additional RI in sympatry | **0.2533** | **0.4764** | -0.1162 | 0.1355 |
| $\beta_{int}$ | increase in slope for sympatry | **-0.3209** | **-0.0593** | **0.1326** | **0.5247** |

**Figure 1.**
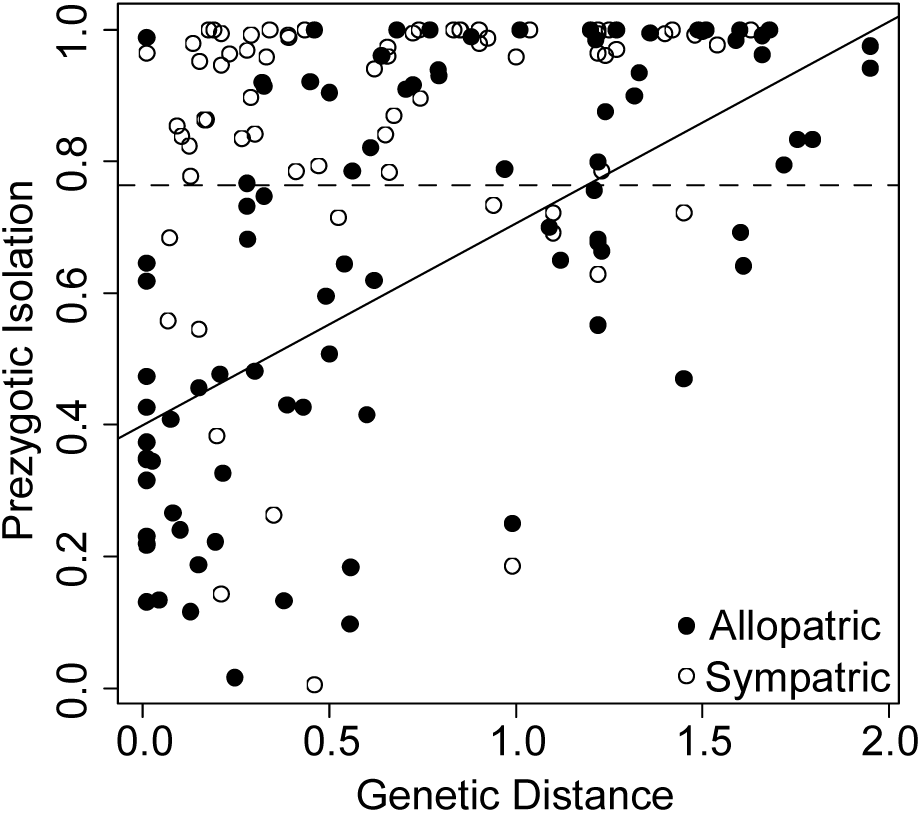
The relationship between genetic distance and prezygotic isolation for *Drosophila* species pairs that are either allopatric or sympatric, demonstrating that most sympatric species pairs show almost complete isolation, resulting in no relationship between genetic distance and reproductive isolation for the sympatric context. The best fit lines were constructed using the mode of the parameters from the MCMCglmm analysis. The solid line represents allopatric species pairs and the dashed line represents sympatric species pairs.

For post-zygotic isolation the intercept and increase in reproductive isolation in sympatry were not significantly different than zero (*µ* and *γ_sym_* had HPD overlapping zero; Table 1). This indicates that there is little to no postzygotic isolation in recently diverged species regardless of geographical context. The relationship between genetic distance and reproductive isolation (*β_gen.dist_*) was significant and the rate of increase of reproductive isolation with genetic distance was greater in sympatric pairs (*β_int_* = 0.1326, 0.5247). This suggests that reproductive isolation may accumulate more quickly between sympatric pairs of species than allopatric pairs, and the difference increases as divergence time (genetic distance) increases.

#### Bufonidae

Analyses of the Bufonidae data using the two alternative indices of reproductive isolation were qualitatively the same (Table 2), so I only discuss results using the original index of reproductive isolation. The model intercept was significantly different than zero (*µ* =0.2980, 0.6945) and the effect of sympatry was to actually decrease the level of reproductive isolation (*γ_sym_* = −0.2816, - 0.0427) though the overall level of reproductive isolation was still non-zero (the HPD for *µ* + *γ_syrn_*=(0.0164,0.6518)). The relationship between genetic distance and reproductive isolation was quite steep (*β_gen.dist_*=3.6212, 5.8819), and the increased rate of accumulation in sympatric pairs was also significant (*β_int_*= 0.4679, 3.5163). In combination, these coefficients suggest that even though there is little reproductive isolation for very recently diverged sympatric pairs (those separated by small genetic distances), reproductive isolation accumulates more quickly for sympatric pairs than allopatric pairs.

**Table 2.** Summary of coefficients estimated for the analysis of postzygotic reproductive isolation from the Bufonidae data. The original index of reproductive isolation (left) was calculated by Malone and Fotenont (2008) following the procedure of Coyne and Orr (1989) and Presgraves (2002). The new index (right) takes into account that reproductive barriers are sequential following Ramsey et al. (2003) The confidence intervals are for 95% Highest posterior density (HPD) and are significant if they do not include zero (in bold).

| Coefficient | Biological meaning | Postzygotic (Original) |  | Postzygotic (New) |  |
| --- | --- | --- | --- | --- | --- |
|  |  | Lower | Upper | Lower | Upper |
| $\mu$ (intercept) | average RI | **0.2980** | **0.6945** | **0.8398** | **0.9686** |
| $\beta_{gen.dist.}$ | slope relating genetic distance and RI | **3.6212** | **5.8819** | **0.4472** | **1.2592** |
| $\gamma_{sym}$ | additional RI in sympatry | **-0.2816** | **-0.0427** | **-0.0988** | **-0.0105** |
| $\beta_{int}$ | increase in slope for sympatry | **0.4679** | **3.5163** | **0.2665** | **1.3992** |

### Silene

Sympatry had no effect on the baseline prezygotic reproductive isolation (*γ_sym_*=-03643, 0.1510) or on the rate of accumulation of reproductive isolation (*β_int_*=-1.5671, 3.5080), which is consistent with the original results from Moyle et al (2004). There was a significant relationship between genetic distance and reproductive isolation (*β_gen.dist_*), which did not differ between sympatry and allopatry (Table 3). The lack of allopatric pairs for the postzygotic and total reproductive isolation measurements precluded analysis of the effects of geographical context on these measures (though the general relationship between reproductive isolation and genetic distance was positive and significant consistent with (Moyle et al. 2004); data not shown).

**Table 3.**
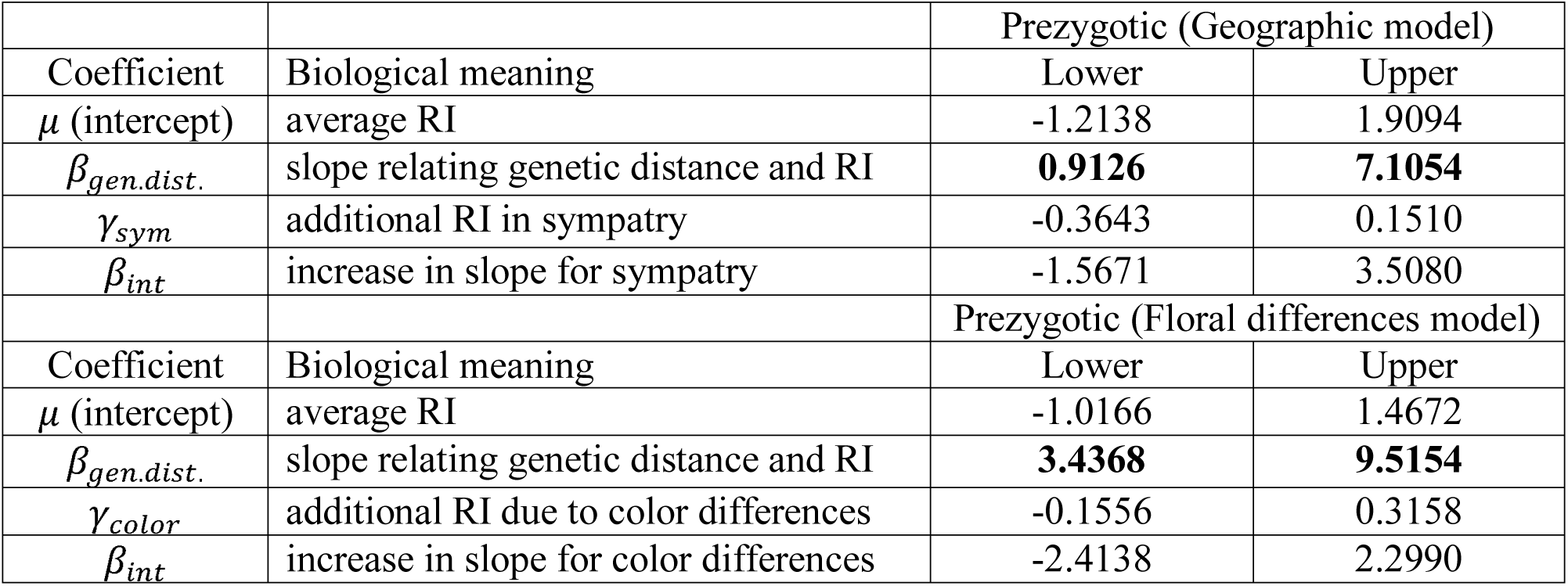
Summary of coefficients estimated for the analysis of prezygotic reproductive isolation when considering geographical context (left) or floral divergence (right) from the *Silene* data. The confidence intervals are for 95% Highest posterior density (HPD) and are significant if they do not include zero (in bold).

|  |  | Prezygotic (Geographic model) |  |
| --- | --- | --- | --- |
| Coefficient | Biological meaning | Lower | Upper |
| $\mu$ (intercept) | average RI | -1.2138 | 1.9094 |
| $\beta_{gen.dist.}$ | slope relating genetic distance and RI | **0.9126** | **7.1054** |
| $\gamma_{sym}$ | additional RI in sympatry | -0.3643 | 0.1510 |
| $\beta_{int}$ | increase in slope for sympatry | -1.5671 | 3.5080 |
|  |  | Prezygotic (Floral differences model) |  |
| Coefficient | Biological meaning | Lower | Upper |
| $\mu$ (intercept) | average RI | -1.0166 | 1.4672 |
| $\beta_{gen.dist.}$ | slope relating genetic distance and RI | **3.4368** | **9.5154** |
| $\gamma_{color}$ | additional RI due to color differences | -0.1556 | 0.3158 |
| $\beta_{int}$ | increase in slope for color differences | -2.4138 | 2.2990 |

For all three measures of reproductive isolation, floral color differences did not increase reproductive isolation. Regardless of the cross category (crosses between species that differed in floral color vs. between species that had the same color) there was a large amount of variation in reproductive isolation with some pairs having little to no isolation, and other pairs having substantial isolation (Figure 2). Thus, there was no effect of floral color on the average levels of reproductive isolation or on the accumulation of reproductive isolation over time, and the only significant effect was genetic distance on reproductive isolation (Table 3; prezygotic model shown).

**Figure 2.**
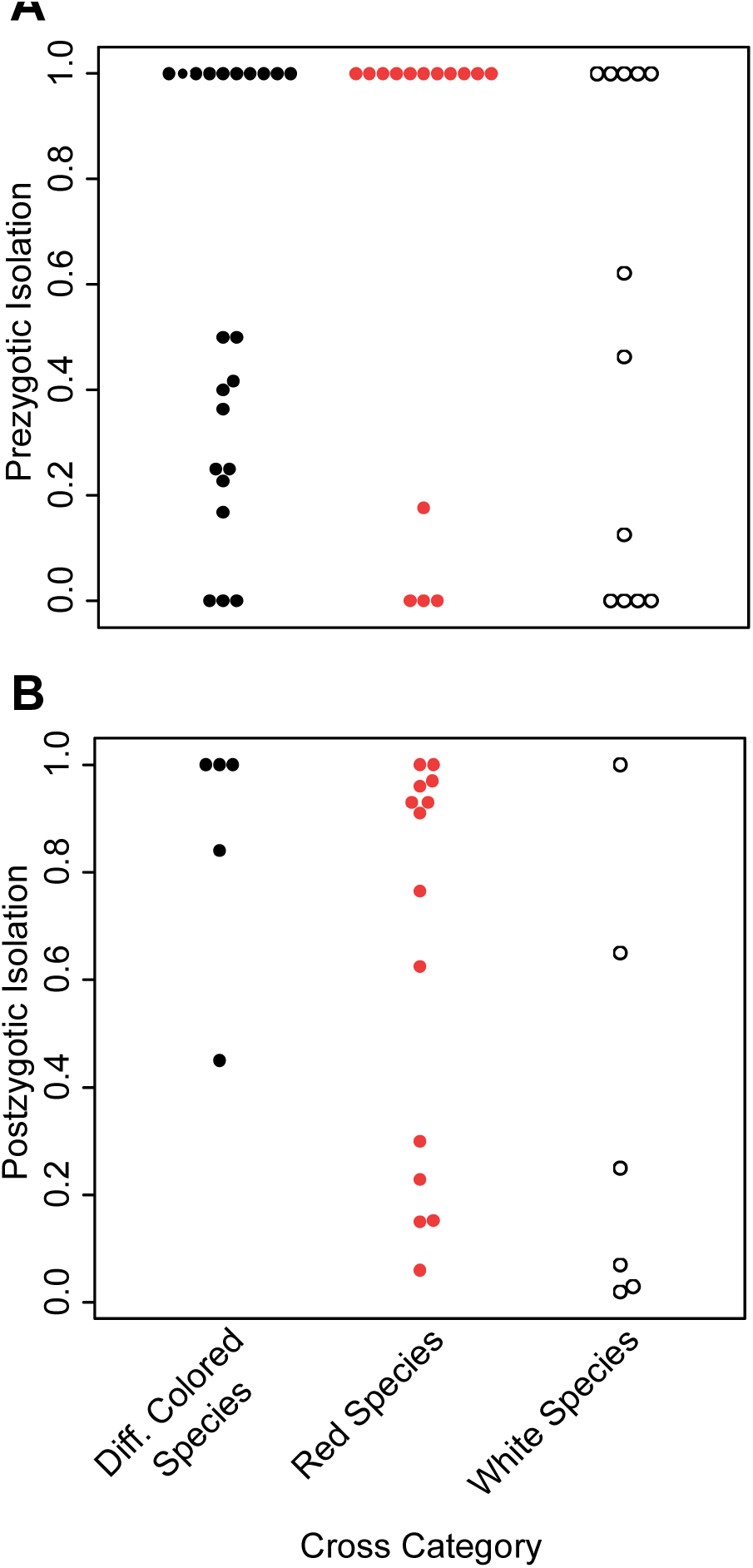
Variation in the level of reproductive isolation between *Silene* species pairs that either had different floral colors or shared floral colors (separated into pairs where both species were either white or red) for both A) prezygotic and B) postzygotic isolation demonstrating no effect of floral differences on reproductive isolation.

### Nolana

When genetic distance, geographic distance, and measures of flower size differences were considered jointly, the only significant predictor of reproductive isolation was geographic distance (*β_geo.dist_*=00001, 0.0004; note, the small coefficient is due to the scale of reproductive isolation/kilometers), signifying that there is more reproductive isolation between pairs of species that are more geographically distant. This is consistent with the original study in the sense that only geographical distance was significant using the Mantel test (Jewell et al. 2012). The analysis using the genetic distance matrix also indicated that geographic distance was the only significant predictor of reproductive isolation (Table 4) and the coefficients were similar to the previous analysis. The main difference was there was no significant intercept (*μ*=(-1.0166, 1.4672)). This is likely caused by the differences in inferred relatedness between the phylogeny and the genetic distance matrix (Supplemental Fig. 2), as this was the only difference in the two models. In a phylogeny relatedness is based on shared ancestry, whereas a distance matrix includes all nucleotide changes without context of whether they are shared with other taxa or phylogenetically informative. The result of using the genetic distance matrix to infer relatedness may have been to infer that more closely related species had little reproductive isolation, so the intercept was not significantly different than zero.

**Table 4.** Summary of coefficients estimated for the analysis of total reproductive isolation when using the phylogeny (left) or genetic distance matrix (right) from the *Nolana* data. The confidence intervals are for 95% Highest posterior density (HPD) and are significant if they do not include zero (in bold).

|  |  | Total reproductive isolation (phylogenetic matrix) |  | Total reproductive isolation (genetic distance matrix) |  |
| --- | --- | --- | --- | --- | --- |
| Coefficient | Biological meaning | Lower | Upper | Lower | Upper |
| $\mu$ (intercept) | average RI | **0.1166** | **0.6249** | -0.3126 | 0.4921 |
| $\beta_{gen.dist.}$ | slope relating genetic distance to RI | -0.3149 | 0.4861 | -0.4088 | 0.3945 |
| $\beta_{geo.dist.}$ | slope relating geographic distance (km) to RI | **0.0001** | **0.0004** | **0.0001** | **0.0004** |
| $\beta_{corolla}$ | slope relating differences in corolla diameter to RI | -0.0071 | 0.0140 | -0.0106 | 0.0115 |

## Discussion

Understanding how reproductive isolation accumulates over time in different geographic contexts (allopatry vs. sympatry) or according to specific trait differences requires analyzing the interaction between genetic distance and these factors and their joint effects on reproductive isolation. These comparisons cannot be made using PICs or Mantel tests, so in many classical studies the effects of these factors are instead indirectly compared (Coyne and Orr 1989, 1997; Moyle et al. 2004; Jewell et al. 2012). Using a phylogenetic mixed model framework, and classic data sets on species ability to cross with one another, I formally tested the interactions among these effects on the evolution of reproductive isolation. I explicitly examined interactions between categorical and continuous variables and accounted for correlations between predictor variables, something that is not possible in the standard PICs framework (Burt 1989; Garland et al. 1992) and difficult to implement using Mantel tests when there are more than two levels of the categorical variable. Mantel tests only evaluate the correlation between two variables but not average effects (i.e. intercepts), which must be tested with other methods, whereas estimates of mean differences between two groups are simultaneously estimated using a mixed model approach.

In addition, in the mixed modeling framework the correlation between predictor variables can be modelled and accounted for through specific variance matrices, which enables several predictors to be tested simultaneously where multicollinearity may have previously limited inferences (as demonstrated in the analysis of *Nolana*). In comparison, current approaches to evaluating the effect of multiple variables on reproductive isolation have clear limitations. Partial Mantel tests do not examine multiple effects simultaneously, but instead evaluate what additional variance a variable may explain after accounting for variance due to other variables (Smouse et al. 1986). Although multiple variables can be accommodated in a standard regression using PICs, it is be difficult to determine the significance/validity of coefficients when the analysis involves correlated independent variables because multicollinearity causes spurious variance inflation (Mundry 2014). This is especially troubling, given that the nature of the data used in these studies will often contain correlations between the variables that may explain the accumulation of reproductive isolation.

The phylogenetic mixed effect model is also flexible in that relationships between taxa can be conveyed either through a phylogenetic relatedness matrix or a genetic distance matrix. In contrast, often the use of PICs vs. Mantel test reflects the scale of relatedness that researchers are examining: PICs typically examine interspecific data using phylogenies while Mantel tests can focus on intraspecific data using genetic distance matrices. The mixed model approach is a natural way to apply these analyses across different phylogenetic scales.

### New insights from phylogenetic mixed model framework

Given the advantages of the phylogenetic mixed model framework I discuss the inferences from the hypotheses I examined, both in the context of how they compared to the original results and what additional inferences can be made.

For the *Drosophila* and Bufonidae analyses I specifically tested whether geographic context (allopatry or sympatry) influences the rate of accumulation of reproductive isolation. These data were both originally analyzed using linear regression (similar to PICS), however the difference in the rate of accumulation (represented by the slope relating genetic distance to reproductive isolation) could not be tested directly because of the inability to calculate contrasts for binary variables and therefore the interaction between geographic context and genetic distance could not be evaluated. Using the phylogenetic mixed model enabled direct tests of the differences in the rate of isolation accumulation, and doing so produced new insights into these previously analyzed datasets. In particular, I found that in both *Drosophila* and Bufonidae there was an increase in the rate of accumulation of postzygotic reproductive isolation in sympatric species pairs compared to allopatric species pairs. This result contrasts with the analysis by Coyne and Orr (1997) and likely reflects the fact that Coyne and Orr (1997) used a subset of the data that only included recently diverged species (D<0.25), whereas analyses here included the entire dataset. Other studies that have included data from both sympatric and allopatric pairs in Lepidoptera (Presgraves 2002) and birds (Price and Bouvier 2002) failed to find differences between sympatry and allopatry in the level of reproductive isolation, although the inferences were based on informal analyses.

For prezygotic isolation, the presence of stronger prezygotic isolation among closely related species pairs in sympatry is often assumed to be a product of reinforcement (Yukilevich 2012), but reinforcement is an unlikely force directly contributing to increased accumulation of postzygotic isolation in sympatry (Servedio 2000; Servedio and Saetre 2003). An alternative hypothesis, called the Templeton Effect or differential fusion effect, proposes that strong reproductive isolation in sympatry is a consequence of a systematic bias among species pairs that are able to maintain their integrity in sympatry, whereby weakly isolated species fail to maintain species boundaries (and therefore undergo species collapse), leaving only species pairs with strong reproductive isolation. This hypothesis has previously been proposed to explain patterns of strong prezygotic isolation in sympatry (Templeton 1981; Noor 1999; Yukilevich 2012), however (unlike reinforcement) the effect may be equally applicable to postzygotic isolation. The Templeton/differential fusion effect may be reinterpreted and applied to postzygotic isolation such that the level of postzygotic isolation that exists between species will determine the likelihood of species coexistence. This is because sexual isolation alone is often not strong enough to maintain species barriers (Lande 1981; Payne and Krakauer 1997; Servedio and Burger 2014), and reinforcement is sensitive to gene flow. If the strength of postzygotic isolation between sympatric lineages drives reinforcement and ultimately the strength of prezygotic isolation (Servedio 2000; Servedio and Saetre 2003), we should observe stronger postzygotic isolation in sympatry (as observed for both the *Drosophila* and Bufonidae data sets) and a correlation between postzygotic and prezygotic isolation in sympatry. Interestingly, postzygotic and prezygotic isolation are correlated in sympatry in the *Drosophila* dataset (Yukelivich 2012).

The mixed model approach can also be used to evaluate the effect of categorical trait variation on the strength and accumulation of reproductive isolation, to address additional mechanistic questions. In my analyses of whether floral divergence can contribute to post-mating and postzygotic reproductive isolation, independent of effects on pollinator visitation, I found no support in *Silene* for the hypothesis that floral divergence could also influence these reproductive isolation phenotypes through pleiotropic effects (Haak et al. 2014). Specifically, floral divergence in the form of flower color differences did not contribute to either prezygotic isolation (most likely via pollen-pistil interactions) or postzygotic isolation (F1 pollen sterility). It is possible that the pleiotropic effects of floral traits on reproductive isolation could manifest in a different traits (seed sterility, F1 germination/viability) or floral divergence could contribute to extrinsic post zygotic isolation (Ramsey et al. 2003; Lowry et al. 2008), all of which were not captured in this analysis. Alternatively (or in addition), differentiation in floral traits other than color might be more important in this context. In *Silene*, for example, there is some evidence that floral traits other than red vs. white flower color, including flower display height and orientation (Brothers and Atwell 2014, Fenster et al. 2015) and floral scent (Waelti et al. 2008, Castillo et al. 2014) may contribute to reproductive isolation via pollinator visitation.

Trait variation like floral variation might also be correlated with other factors that contribute to reproductive isolation. For example, if floral shape has phylogenetic signal then floral divergence would be correlated with genetic distance; similarly, if there is selection for different flower sizes in different environments, floral divergence would be associated with geographical distance. Under these scenarios floral divergence might seem to be contributing to reproductive isolation, merely because it shares a main driving factors with the accumulation of isolation, so it is important to be able to distinguish these potential mechanisms. In the original *Nolana* analysis (Jewell et al. 2012), floral, geographical, and genetic distance measures were all observed to be correlated. In the reanalysis here, I did not observe an independent effect of floral divergence (floral size difference) on reproductive isolation. Indeed, I was able to explicitly rule out the possibility of floral changes contributed to reproductive isolation while simultaneously testing the effects of genetic distance and geographic distance, and found geographical distance alone contributed to reproductive isolation in this system. This result might reflect both reduced gene flow between geographically distant species and/or differences in habitats, both of which could contribute to reproductive isolation. Regardless, it is clear that, differences in flower size do not appear to relay into pleiotropic effects on reproductive isolation in this system, as has been hypothesized for differences in traits associated with pollinator preference (Haak 2014).

### Conclusion

The phylogenetic mixed model framework utilized in this study remedies difficulties for Mantel tests and PICs in testing hypothesis about factors contributing to the evolution of reproductive isolation. To demonstrate the utility of this framework, I performed several analyses to evaluate the roles of categorical geographic and trait variation, and quantitative divergence measures, on the accumulation and strength of isolation in four published datasets. I tested the role of geography in the evolution of reproductive isolation and was able to show that reproductive isolation accumulates more quickly in sympatry not only for prezygotic isolation but also for postzygotic isolation. In the datasets examined, floral traits did not contribute to the pattern or strength of reproductive isolation measures included in the original studies. This framework can enable future studies to test complex hypothesis, test the effects of multiple variables simultaneously (even if they are correlated), and use a generalized framework to examine reproductive isolation between species or at the intraspecies level.

## Supplementary Information

**Supplemental Figure 1.**
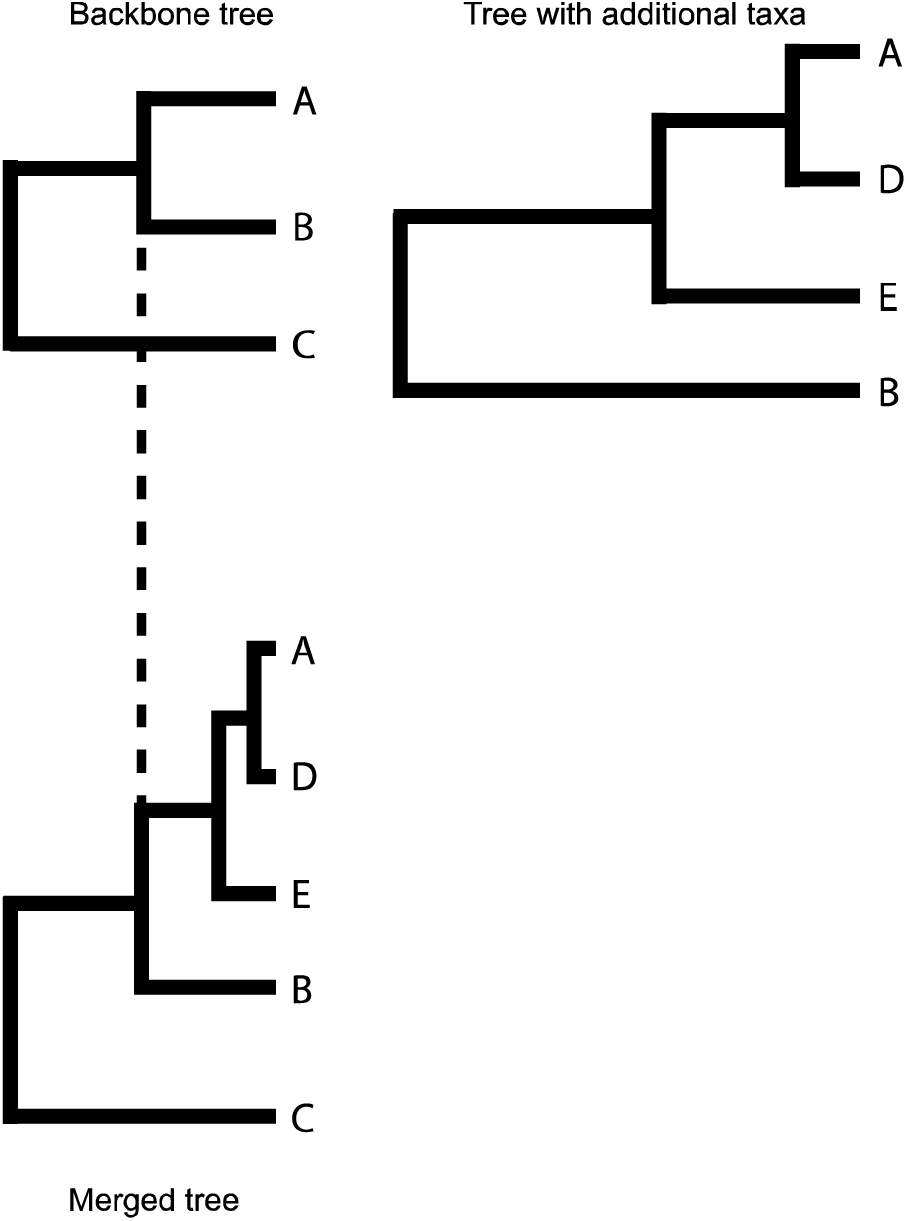
A visual representation of how more detailed information about a species group found in one tree can be integrated into a larger phylogeny by transforming branch lengths. In this example the relationships between species A, D, E, and B are scaled to the same length that occurred in the backbone tree (for the clade consisting of species A and B).

**Supplemental Figure 2.**
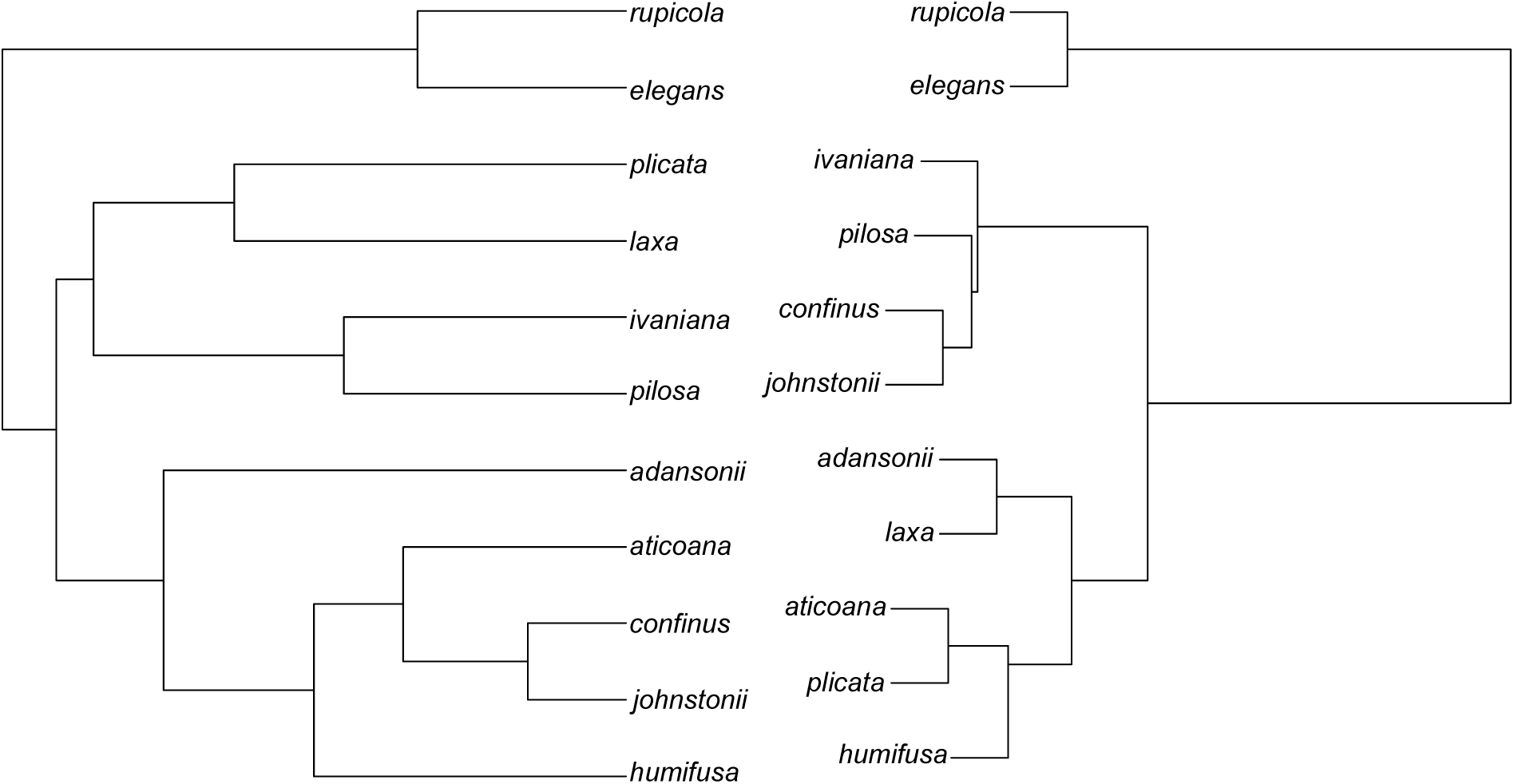
Comparison of relationships in the genus *Nolana* when either a phylogeny made from several loci or matrix of pairwise genetic distances are used. The topology on the left represents the maximum likelihood tree based on sequence from the *ADH2, atpB, ndhF, psbA-trnH, rps16* genes. The topology on the right was generated using Ward's hierarchical clustering method for pairwise genetic distances reported in Jewell et al (2012).

## Literature Cited

Bivand, R., T. Keitt, and B. Rowlingson. 2013. rgdal: Bindings for the geospatial data abstraction library. R package version 0.8-6.

Bivand, R., and N. Lewin-Koh. 2013. maptools: Tools for reading and handling spatial objects. R package version 0.8-23.

Brooks, S. P., and A. Gelman. 1998. General methods for monitoring convergence of iterative simulations. J. Comput. Graph. Stat. 7:434–455.

Brothers, A. N., and J. W. Atwell. 2014. The role of pollinator-mediated selection in the divergence of floral traits between two closely related plant species. Int. J. Plant Sci. 175:287–295

Burt, A. 1989. Comparative methods using phylogenetically independent contrasts. *in* P.H. Harvey and L. Partridge eds, Oxford Surveys in Evolutionary Biology 6:33–53.

Castillo, D. M., A. A. R. Kula, S. Dötterl, M. R. Dudash, and C. B. Fenster. 2014. Invasive *Silene latifolia* may benefit froma native pollinating seed predator, *Hadena ectypa*, in North America. Int. J. Plant Sci. 175:80–91.

Coyne, J. A., and H. A. Orr. 1989. Patterns of speciation in Drosophila. Evolution 43:362–381.

Coyne, J. A., and H. A. Orr. 1997. “Patterns of speciation in *Drosophila*” revisited. Evolution 51:295–303.

Felsenstein, J. 1985. Phylogenies and the comparative method. Am. Nat. 125:1–15.

Fenster, C. B., W. S. Armbruster, P. Wilson, M. R. Dudash, and J. D. Thomson. 2004. Pollination syndromes and floral specialization. Annu. Rev. Ecol. Evol. Syst. 35: 375–403.

Fenster, C. B., R. J. Reynolds, C. W. Williams, R. Makowsky, and M. R. Dudash. 2015. Quantifying hummingbird preference for floral trait combinations: The role of selection on trait interactions in the evolution of pollination syndromes. Evolution 69:1113–1127.

Fitzpatrick, B. M., and M. Turelli. 2006. The geography of mammalian speciation: mixed signals from phylogenies and range maps. Evolution 60:601–615.

Funk, D. J., P. Nosil, and W. J. Etges. 2006. Ecological divergence exhibits consistently positive associations with reproductive isolation across disparate taxa. Proc. Natl. Acad. Sci. U.S.A. 103:3209–3213.

Garland, T., P. H. Harvey, and A. R. Ives. 1992. Procedures for the analysis of comparative data using phylogenetically independent contrasts. Syst. Biol. 4:18–32.

Gelman, A., and D. Rubin. 1992. Inference from iterative simulation using multiple sequences. Stat. Sci. 7:457–511.

Grafen, A. 1989. The phylogenetic regression. Philos. Trans. R. Soc. Lond. B Biol. Sci. 326:119–157.

Grant, V. 1992. Floral isolation between ornithophilous and sphingophilous species of *Ipomopsis* and *Aquilegia*. Proc. Natl. Acad. Sci. U.S.A. 89:11828–11831.

Haak, D. C., J. L. Kostyun, and L. C. Moyle. 2014. Merging ecology and genomics to dissect diversity in wild tomatoes and their relatives. *in* C.R. Landry and N. Aubin-Horth eds, Ecological Genomics: Ecology and the Evolution of Genes and Genomes 781:273–298.

Hadfield, J. D. 2010. MCMC methods for multi-response generalized linear mixed models: The MCMCglmm R package. J. Stat. Softw. 33.

Hadfield, J. D., B. R. Krasnov, R. Poulin, and S. Nakagawa. 2014. A tale of two phylogenies: Comparative analyses of ecological interactions. Am. Nat. 183:174–187.

Hadfield, J. D., and S. Nakagawa. 2010. General quantitative genetic methods for comparative biology: phylogenies, taxonomies and multi-trait models for continuous and categorical characters. J. Evol. Biol. 23:494–508.

Harmon, L. J., and R. E. Glor. 2010. Poor statistical performance of the Mantel test in phylogenetic comparative analyses. Evolution 64:2173–2178.

Hodges, S. A., and M. L. Arnold. 1994. Floral and ecological isolation between *Aquilegia formosa* and *Aquilegia pubescens*. Proc. Natl. Acad. Sci. U.S.A. 91:2493–2496.

IUCN. 2014. The IUCN Red List of Threatened Species 2014. Available at www.iucnredlist.org. Accessed October 21, 2014.

Jenkins, C., and S. Keller. 2011. A phylogenetic comparative study of preadaptation for invasiveness in the genus *Silene* (Caryophyllaceae). Biol. Invasions 13:1471–1486.

Jewell, C., A. D. Papineau, R. Frefre, and L. C. Moyle. 2012. Patterns fo reproductive isolation in *Nolana* (Chilean bellflower). Evolution 66:2628–2636.

Kay, K. M., and R. D. Sargent. 2009. The role of animal pollination in plant speciation: integrating ecology, geography, and genetics. Annu. Rev. Ecol. Evol. Syst. 40:637–656.

Lande, R. 1981. Models of speciation by sexual selection on polygenic traits. Proc. Natl. Acad.Sci. USA 78:3721–3725.

Lee, C. B., L. E. Page, B. A. McClure, and T. P. Holtsford. 2008. Post-pollination hybridization barriers in *Nicotiana* section Alatae. Sex. Plant Reprod. 21:183–195.

Legendre, P. 2000. Comparison of permutation methods for the partial correlation and partial Mantel tests. J. Stat. Comp. Sim. 67:37–73.

Legendre, P., and M. Fortin. 2010. Comparison of the Mantel test and alternative approaches for detecting complex multivariate relationships in the spatial analysis of genetic data. Mol. Ecol. Res. 10:831–844.

Lowry, D. B., J. L. Modliszewski, K. M. Wright, C. A. Wu, and J. H. Willis. 2008. The strength and genetic basis of reproductive isolating barriers in flowering plants. Philos. Trans. R.Soc. Lond. B Biol. Sci. 363:3009–3021.

Malone, J. H., and B. E. Fontenot. 2008. Patterns of reproductive isolation in toads. PLoS One 3:11.

Mantel, N. A. 1967. The detection of disease clustering and a generalized regression approach. Cancer Res. 27:209–220.

Meiners, J., and T. Winkelmann. 2012. Evaluation of reproductive barriers and realisation of interspecific hybridisations depending on genetic distances between species in the genus *Helleborus*. Plant Biol. 14:576–585.

Mendelson, T. C. 2003. Sexual isolation evolves faster than hybrid inviability in a diverse and sexually dimorphic genus of fish (Percidae: Etheostoma). Evolution 57:317–327.

Morales-Hojas, R., and J. Vieira. 2012. Phylogenetic patterns of geographical and ecological diversification in the subgenus *Drosophila*. PLoS One 7.

Moyle, L. C., M. S. Olson, and P. Tiffin. 2004. Patterns of reproductive isolation in three angiosperm genera. Evolution 58:1195–1208.

Mundry, R. 2014. Statistical issues and assumptions of phylogenetic generalized least squares. *in* L. Z. Garamszegi ed, Modern Phelogenetic Comparative Methods and Their Application in Evolutionary Biology. Springer-Verlag, Berlin.

Nei, M. 1972. Genetic distance between populations. Am. Nat. 106:283–292.

Nei, M., F. Tajima, and Y. Tateno. 1983. Accuracy of estimated phylogenetic trees from molecular data. J. Mol.Evol. 19:153–170.

Noor, M. A. F. 1999. Reinforcement and other consequences of sympatry. Heredity 83:503–508.

Nosil, P. 2013. Degree of sympatry affects reinforcement in *Drosophila*. Evolution 67:868–872.

Payne, J.H., and D.C. Krakauer. 1997. Sexual selection, space and speciation. Evolution 51:1–9.

Plummer, M., N. Best, K. Cowles, and K. Vines. 2006. CODA: Convergence diagnosis and output analysis for MCMC. R News 6.

Presgraves, D. C. 2002. Patterns of postzygotic isolation in Lepidoptera. Evolution 56:1168–1183.

Price, T. D., and M. M. Bouvier. 2002. The evolution of F1 postzygotic incompatabilities in birds. Evolution 56:2083–2089.

Ramsey, J., H. D. Bradshaw, and D. W. Schemske. 2003. Components of reproductive isolation between the monkeyflowers *Mimulus lewisii* and *M. cardinalis* (Phrymaceae). Evolution:1520–1534.

Servedio, M.R. 2000. Reinforcement and the genetics of nonrandom mating. Evolution 54:21–29

Servedio, M. R., and R. Burger. 2014. The counterintuitive role of sexual selection in species maintenance and speciation. Proc. Natl. Acad. Sci. USA 111:8113–8118.

Servedio, M.R. and G.-P. Sætre. 2003. Speciation as a positive feedback loop between post-and prezygotic barriers to gene flow. Proc. Roy. Soc. Lond. B 270:1473–1479.

Schemske, D. W., and H. D. Bradshaw. 1999. Pollinator preference and the evolution of floral traits in monkeyflowers (*Mimulus*). Proc. Natl. Acad. Sci. U.S.A. 96:11910–11915.

Schnute, J. T., N. Boers, and R. Haigh. 2012. PBSmapping: Mapping fisheries data and spatial analysis tools. R package version 2.65.40.

Slatkin, M., M. Veuille, and J. Felsenstein. 2002. Contrasts for a within-species comparative method. *in* M. Slatkin and M. Veuille eds, Modern developments in theoretical population genetics. Oxford University Press, Oxford, UK.

Smaczniak, C., R. G. H. Immink, G. C. Angenent, and K. Kaufmann. 2012. Developmental and evolutionary diversity of plant MADS-domain factors: insights from recent studies. Development 139:3081–3098.

Smith, R. A., and M. D. Rausher. 2007. Close clustering of anthers and stigma in *Ipomoea hederacea* enhances prezygotic isolation from *Ipomoea purpurea*. New Phytol. 173:641–647.

Smouse, P. E., J. C. Long, and R. R. Sokal. 1986. Multiple regression and correlation extensions of the Mantel test of matrix correspondence. Syst. Zool. 35:627–632.

Stamatakis, A. 2014. RAxML Version 8: A tool for phylogenetic analysis and post-analysis of large phylogenies. Bioinformatics doi: 10.1093/bioinformatics/btu033.

Stone, G. N., S. Nee, and J. Felsenstein. 2011. Controlling for non-independence in comparative analysis of patterns across populations within species. Philos. Trans. R. Soc. Lond. B Biol. Sci. 366:1410–1424.

Templeton, A. R. 1981. Mechanisms of speciation a population genetic approach. Annu. Rev. Ecol. Evol. Syst. 12:23–48.

Turelli, M., J. R. Lipkowitz, and Y. Brandvain. 2014. On the Coyne and Orr-igin of species: effects of intrinsic postzygotic isolation, ecological differentiation, X chromosome size, and sympatry on Drosophila speciation. Evolution 68:1176–1187.

van der Linde, K., D. Houle, G. S. Spicer, and S. J. Steppan. 2010. A supermatrix-based molecular phylogeny of the family *Drosophilidae*. Genet. Res. 92:25–38.

Venables, W. N., and B.D Ripley. 2002. Modern Applied Statistics with S. 4th ed. Springer, New York.

Waelti, M. O., J. K. Muhlemann, A. Widmer, and F. P. Schiestl. 2008. Floral odour and reproductive isolation in two species of *Silene*. J. of Evol. Biol. 21:111–121.

Wang, I. J. 2013. Examining the full effects of landscape heterogeneity on spatial genetic variation: A multiple matrix regression approach for quantifying geographic and ecological isolation. Evolution 67:3403–3411.

Weir, B. S., and C. C. Cockerham. 1984. Estimating F-Statistics for the analysis of population structure. Evolution 38:1358–1370.

Yukilevich, R. 2012. Asymmetrical patterns of speciation uniquely support reinforcement in *Drosophila*. Evolution 66:1430–1446.

